# Delta C2 Domain β1-2 loop contributes to robust Notch signalling

**DOI:** 10.1101/2021.02.16.431397

**Authors:** Torcato Martins, Yao Meng, Boguslawa Korona, Richard Suckling, Steven Johnson, Penny Handford, Susan M. Lea, Sarah Bray

**Affiliations:** Department of Physiology Development and Neuroscience, University of Cambridge, Downing Street, Cambridge CB2 3DY, UK; Department of Biochemistry, University of Oxford, Oxford, UK; Sir William Dunn School of Pathology, University of Oxford, Oxford, UK

**Keywords:** Delta, *Drosophila*, ligands, C2 domain, membrane composition, CRISPR/Cas9, signalling

## Abstract

Accurate Notch signalling is critical for organism development and homeostasis. Fine-tuning of Notch-ligand interactions have substantial impact on signalling-outputs. Recent structural studies identified a conserved N-terminal C2 domain in human Notch ligands which conferred phospholipid binding in vitro. Here we show that Drosophila ligands adopt the same C2 domain structure with analogous variations in the loop regions, including the so-called β1-2 loop that has been associated with phospholipid binding. Mutations in the β 1-2 loop of Delta C2 domain retain Notch binding but have impaired ability to interact with phospholipids in vitro. To investigate its role in vivo we deleted five residues within the β 1-2 loop of endogenous Delta by CRISPR/Cas9 gene editing. Strikingly, this change compromised ligand function. The modified Delta enhanced phenotypes produced by Delta loss of function alleles and suppressed that of Notch alleles. As the modified protein was present on the cell surface in normal amounts, these results argue that C2 domain phospholipid-binding is necessary for robust signalling in vivo where the balance of cis and trans ligand-receptor interactions is finely tuned.

## INTRODUCTION

The Notch signalling pathway is highly conserved and plays key roles in many aspects of development and homeostasis (Bray, 2016). Aberrant Notch signalling results in a number of inherited diseases and is associated with various cancers and other acquired disorders (Mašek & Andersson, 2017; Nowell & Radtke, 2017; Siebel & Lendahl, 2017; Monticone & Miele, 2021). As both the Notch receptors and the ligands are single‐pass type I transmembrane proteins, signalling is initiated by direct protein–protein contact between adjacent cells, which may occur in some instances via long cell processes such as cytonemes (Boukhatmi *et al*, 2020; Cohen *et al*, 2010; Hunter *et al*, 2019; De Joussineau *et al*, 2003; Huang & Kornberg, 2015). Canonical Notch signalling involves a simple cascade, whereby ligand binding induces successive cleavages to release the Notch intracellular domain (NICD) which translocates to the nucleus and directly regulates gene expression with its binding partners (Bray, 2016; Kovall *et al*, 2017; Kovall & Blacklow, 2010; Kovall, 2008). One challenge is to understand how this simple core mechanism is modulated to ensure appropriate spatio-temporal regulation of the pathway. Mechanisms that fine-tune the ligand receptor interactions are likely to make important contributions.

All Notch ligands have a similar architecture, with an extracellular domain consisting of multiple (7, 8, or 16) epidermal growth factor (EGF) repeats, a so-called Delta/Serrate/Lag‐2 (DSL) domain and a highly conserved N-terminal region (Kopan & Ilagan, 2009; D’Souza *et al*, 2008; Bray, 2006; Kovall & Blacklow, 2010). Receptor binding involves the N-terminal portion including the DSL and N-terminal domains (Luca *et al*, 2015, 2017; Cordle *et al*, 2008). Structural studies of the N-terminal region from human Delta and Jagged ligands revealed that it adopts a conformation characteristic of a phospholipid-binding C2 domain (Chillakuri *et al*, 2013; Kershaw *et al*, 2015). In agreement, these domains interact with phospholipid-containing liposomes *in vitro*, and exhibit ligand-specific preferences for liposomes of different composition (Suckling *et al*, 2017). Comparisons between mammalian Jagged and Delta type ligands revealed a diversity in the structures of the loops at the apex of the C2 domain which are implicated in membrane recognition in other C2 domain proteins (Suckling *et al*, 2017). A subset of missense mutations, which affect these loops in Jagged1 are associated with extrahepatic biliary atresia (EHBA)(Kohsaka *et al*, 2002). Purified EHBA variants show reduced Notch activation in reporter cell assays, lead to a reduction in phospholipid binding, but do not alter Notch binding (Suckling *et al*, 2017). The C2 domain may therefore have a role in tuning the activity of the Notch ligands through its lipid-binding properties.

Mutations affecting the single Delta or Serrate (Jagged-like) ligands in *Drosophila* have well characterized consequences on development (e.g. (Heitzler & Simpson, 1991; Thomas *et al*, 1991; Fleming, 1998; Bishop *et al*, 1999; de Celis *et al*, 1996, 1997). Homozygous loss of ligand function leads to lethality but several defects, including wing venation abnormalities, are detected even in *Delta* heterozygotes, which have one normal gene copy (Dexter, 1914; Huppert *et al*, 1997; de Celis *et al*, 1997). As these defects occur when only one allele is mutated, it is evident that patterning is highly sensitive to ligand levels and activity. This therefore provides a powerful context in which to investigate the contributions from the apical C2 domain loops to ligand activity *in vivo*.

As the loop regions of the C2 domains are the most variable, we first set out to solve the structure of the C2 domains from the *Drosophila* Delta and Serrate ligands. This revealed similar prominent β1-2 and β5-6 loops to those in the C2 domain of the mammalian ligands that are thought to be responsible for the interaction with phospholipid head groups (Suckling *et al*, 2017). To test the functional contribution, we focussed on the β1-2 loop in Delta and used CRISPR/Cas9 genome editing to delete 5 amino acids so that we could analyse the impact on Notch activity during development. *In vitro*, such *Dl^Δβ1-2^* mutation(s), resulted in expression of stable protein with altered phospholipid-binding properties. Strikingly, *in vivo* the *Dl^Δβ1-2^* mutation compromised ligand function, exhibiting characteristics of reduced signalling activity. Our data therefore confirm the relevance of C2 domain loops for full ligand activity and, given their ability to confer lipid-binding, suggests that membrane-binding properties are important for robust signalling.

## RESULTS

### Structure and binding properties of the C2 domain of *Drosophila* ligands

To determine whether the *Drosophila* ligands adopt the same arrangement as their mammalian counterparts we solved the structures of the N-terminal region of *Drosophila* Delta and Serrate (Fig. 1) as well as the ligand binding region of *Drosophila* Notch (EGF11-13; Fig. EV1). These were solved using molecular replacement of the individual domains from the human homologues to resolutions between 1.5 and 3.0 Å (Table 1, Fig. 1). When the new *Drosophila* ligand structures were overlaid on their mammalian equivalents, Jagged-1 and DLL4, it was evident that the core domain structure and arrangements of the fly ligands are highly conserved (Fig. 1A-C and Fig. EV1; RMSD 2.5 Å for Delta and 3.1 Å for Serrate) as was the structure and domain arrangement of the Notch receptor ligand binding region (Fig. EV1; RMSD 1.1 Å). The conserved domain arrangement allows us to model the Notch-ligand complex by overlay of the *Drosophila* structures on the earlier structures of the mammalian complexes (Luca *et al*, 2015, 2017) with this leading to no significant clashes between the Notch and ligand coordinates. Notable exceptions to the overall conserved arrangements are the β1-2 and β5-6 loops which exhibit different lengths and folding in the ligands. These highly variable loops protrude apically from the C2 domain core and are positioned far from the Notch-binding interface.

**Figure 1.**
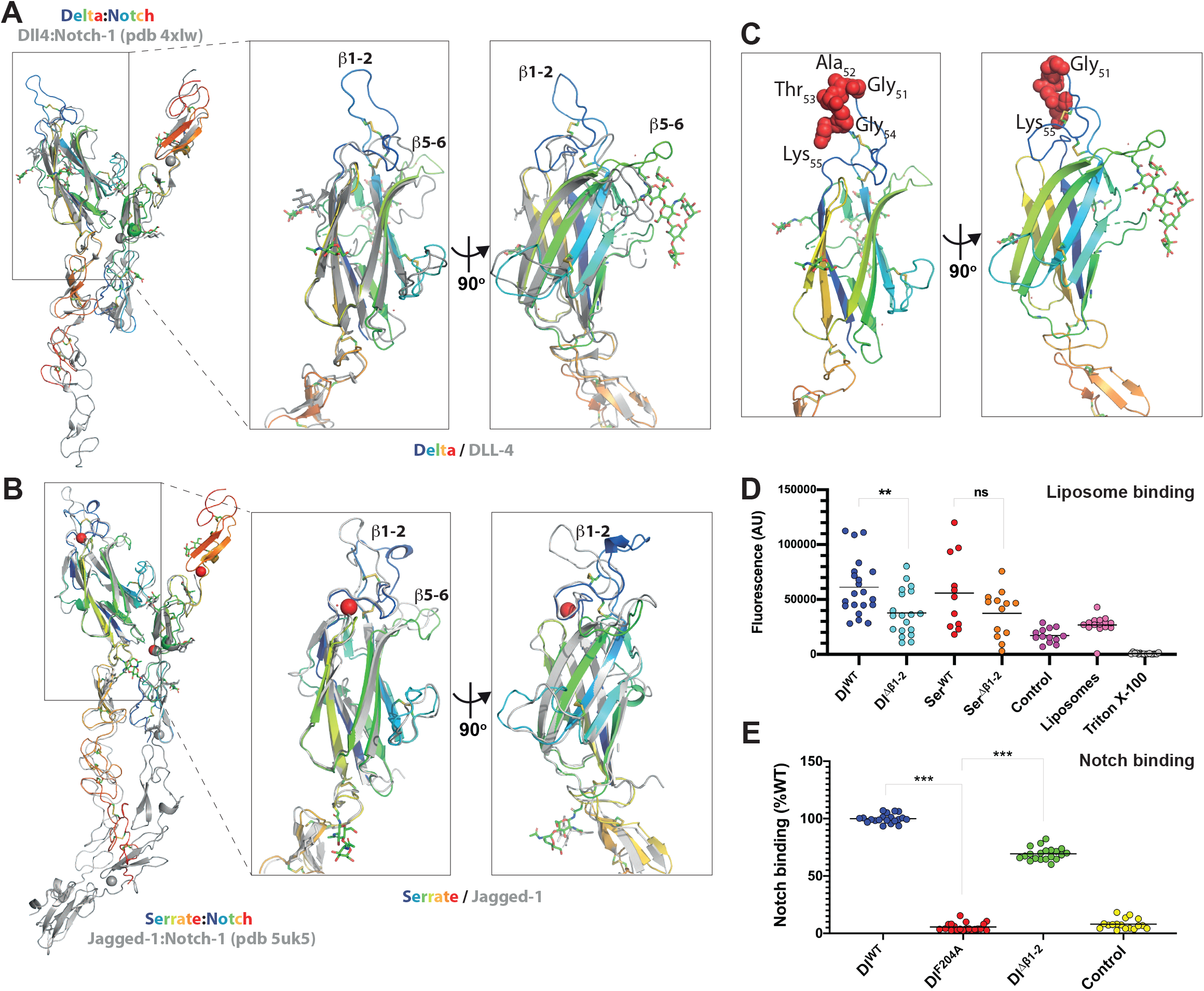
Structure and binding properties of *Drosophila* ligands. **(A,B)** Left panels. The structures of the N-terminal regions from *Drosophila* Delta (**A**) and Serrate (**B**) are shown in a cartoon representation (rainbow coloured from blue at N-terminus to red at C-terminus). These have been overlaid on their mammalian equivalents DLL-4 (**A**) and Jagged-1 (**B**) in the context of their complexes (PDB entries 4xlw and 5uk5 respectively) with Notch-1 (cartoon representation, coloured grey). The structure of isolated *Drosophila* Notch is also depicted in each panel (cartoon, rainbow coloured) superposed on the respective copy of mammalian Notch-1 (cartoon, grey) for each complex. The overlays demonstrate the high degree of conservation in domain structures and arrangements between the *Drosophila* and mammalian homologues. Right panels. A close-up view of the C2 domains of each ligand overlaid with their mammalian equivalent. These demonstrate conservation of overall fold but large differences in the apical loops, particularly in the β1-2 and β5-6 loops. **(C)** Isolated structure of N-terminal Delta, with residues deleted in Δβ1-2 highlighted as red Van Der Waals spheres. **(D,E)** Purified *Drosophila* Delta and Serrate NE3 proteins bind to liposomes **(D)** Binding is reduced in Delta^Δβ1-2^, but not to the Serrate equivalent using liposomes composed of PC:PS:PE-fluoroscein (80:15:5). Notch binding to *Drosophila* Delta NE3 **(E)** WT and Δβ1-2 (Delta^Δβ1-2^) variants both bind to Notch, unlike variant with F204A substitution in DSL domain. Comparisons were performed with a two-tailed unpaired t-test. Values are shown as scattered data points with the dark lines representing the means. ns, no significant difference, \*\**P* < 0.01; \*\*\**P* < 0.0001.

**Table 1.** Data collection and refinement statistics.

|  | **Delta C2-DSL-EGF1**<br>**(7ALK)** | **Notch EGF11-13**<br>**(7ALJ)** | **Serrate C2-DSL-**<br>**EGF1-2 (7ALT)** |
| --- | --- | --- | --- |
| **Wavelength** |  |  |  |
| **Resolution range** | 29.11 - 3.0 (3.107 - 3.0) | 45.2 - 1.523 (1.577 - 1.523) | 45.61 - 2.03 (2.103 - 2.03) |
| **Space group** | P 21 | C 2 | P 21 |
| **Unit cell** | 30.99 86.736 47.558<br>90 94.271 90 | 180.82 31.2858<br>21.7952 90 90.769<br>90 | 70.329 49.402<br>93.123 90 110.249<br>90 |
| **Total reflections** | 16842 (1759) | 61424 (6192) | 126615 (12180) |
| **Unique reflections** | 5024 (520) | 18808 (1771) | 38618 (3814) |
| **Multiplicity** | 3.4 (3.5) | 3.3 (3.2) | 3.3 (3.2) |
| **Completeness (%)** | 99.84 (100.00) | 98.38 (92.19) | 98.89 (98.63) |
| **Mean I/sigma(I)** | 4.55 (0.85) | 7.75 (1.68) | 11.10 (2.28) |
| **Wilson B-factor** | 33.66 | 20.54 | 31.51 |
| **R-merge** | 0.2597 (1.459) | 0.07143 (0.4754) | 0.06213 (0.6113) |
| **R-meas** | 0.3097 (1.73) | 0.08538 (0.5717) | 0.0744 (0.7343) |
| **R-pim** | 0.167 (0.9225) | 0.04621 (0.313) | 0.04049 (0.4025) |
| **CC1/2** | 0.952 (0.483) | 0.996 (0.701) | 0.998 (0.769) |
| **CC\*** | 0.988 (0.807) | 0.999 (0.908) | 1 (0.932) |
| **Reflections used in refinement** | 5024 (520) | 18654 (1770) | 38601 (3808) |
| **Reflections used for R-free** | 287 (33) | 962 (103) | 1996 (185) |
| **R-work** | 0.2407 (0.2909) | 0.2006 (0.3670) | 0.2200 (0.3143) |
| **R-free** | 0.2968 (0.3939) | 0.2407 (0.4334) | 0.2578 (0.3285) |
| **CC(work)** | 0.881 (0.588) | 0.946 (0.796) | 0.950 (0.787) |
| **CC(free)** | 0.829 (0.310) | 0.918 (0.819) | 0.906 (0.770) |
| **Number of non-hydrogen atoms** | 1902 | 1024 | 4255 |
| **macromolecules** | 1801 | 856 | 4079 |
| **ligands** | 99 | 71 | 58 |
| **solvent** | 2 | 97 | 118 |
| **Protein residues** | 239 | 115 | 539 |
| **RMS(bonds)** | 0.003 | 0.017 | 0.005 |
| **RMS(angles)** | 0.58 | 1.40 | 0.81 |
| **Ramachandran favored (%)** | 90.31 | 95.58 | 93.95 |
| **Ramachandran allowed (%)** | 9.69 | 4.42 | 5.67 |
| **Ramachandran outliers (%)** | 0.00 | 0.00 | 0.38 |
| **Rotamer outliers (%)** | 0.50 | 0.00 | 1.78 |
| **Clashscore** | 5.19 | 9.82 | 3.98 |
| **Average B-factor** | 35.61 | 31.83 | 41.24 |
| **macromolecules** | 35.01 | 30.95 | 40.91 |
| **ligands** | 46.88 | 36.00 | 67.50 |
| **solvent** | 19.82 | 36.54 | 39.79 |

Statistics for the highest-resolution shell are shown in parentheses.

Given the structural conservation with the mammalian ligands, it is likely that the *Drosophila* proteins exhibit similar properties. Purified N-terminal fragments (NE3 variants) were therefore used to test the liposome-binding capability of variants in which 5 amino acids were deleted from the β1-2 loop, hereafter referred to as Delta^Δβ1-2^. Using a liposome composition of PC:PS:PE-fluoroscein (80:15:5), we could detect binding of wild-type Delta (Delta^WT^) fragment to the liposomes as seen for mammalian Notch ligands (Fig. 1D). This binding was compromised when the variable β1-2 loop was shortened, resulting in the deletion of residues GATGK; Delta^Δβ1-2^ fragment exhibited a significant reduction in binding compared to that from Delta^WT^. Likewise, the equivalent fragment containing Serrate^Δβ1-2^ loop deletion (removal of residues LRATK) also exhibited reduced binding to liposomes, although to a variable extent that was not reproducibly significant (Fig. 1D). This may be due to differences in the lipid-binding specificities because we have previously noted the heterogeneity of the C2 loops sequences in different ligand families and hypothesised that they may confer different lipid-binding specificities (Suckling *et al*, 2017).

Purified Delta (NE3 fragment) also exhibited robust binding to a fragment of *Drosophila* Notch (dNotch EGF11-13), which contains the core ligand-binding sites (Fig. 1E). This relies on the conventional contact sites because it is abolished by an alanine substitution in the DSL domain which replaces a key receptor-binding residue (F204). In comparison, the variant with the loop deletion, the Delta^Δβ1-2^ fragment, retained Notch binding as predicted from the fact that the loop is positioned far away from the Notch binding interface (Fig. 1A-C).

Together these data demonstrate that the C2 domain structure is conserved between species and that the properties detected in the mammalian ligands are also shared by the *Drosophila* counterparts. The main source of variability is present in the N-terminal apical loops which nevertheless are important for liposome binding in *Drosophila* Delta as in DLL-4 and Jagged-1 from mammals.

### Phenotypes produced by mutations in the ligand β1-2 loop

The β1-2 loop is encoded by a small sequence in exon 2 of *Delta* and in exon 3 of *Serrate*, respectively. In order to study the importance of this loop for Notch signalling, the endogenous exons were replaced by modified exons where the coding sequence of the loops was deleted by CRISPR-mediated homologous recombination. For each of the ligands, two gRNAs were designed to flank the target exon, and the recombination of the modified exon was promoted by a complementary sequence within which the β1-2 loop was replaced by a mutated version (*Dl^Δβ1-2^*; Fig. 2A and EV2A). Successful recombination was identified by the presence of a dsRed marker that was subsequently removed and the mutations were confirmed by sequencing of the exon. As well as generating *Dl^Δβ1-2^* mutations, we also recovered a deletion of the entire exon 2, *Dl^ΔExon2^* which removes a key part of the receptor-binding region and behaves as a null allele (Fig. EV2E).

**Figure 2.**
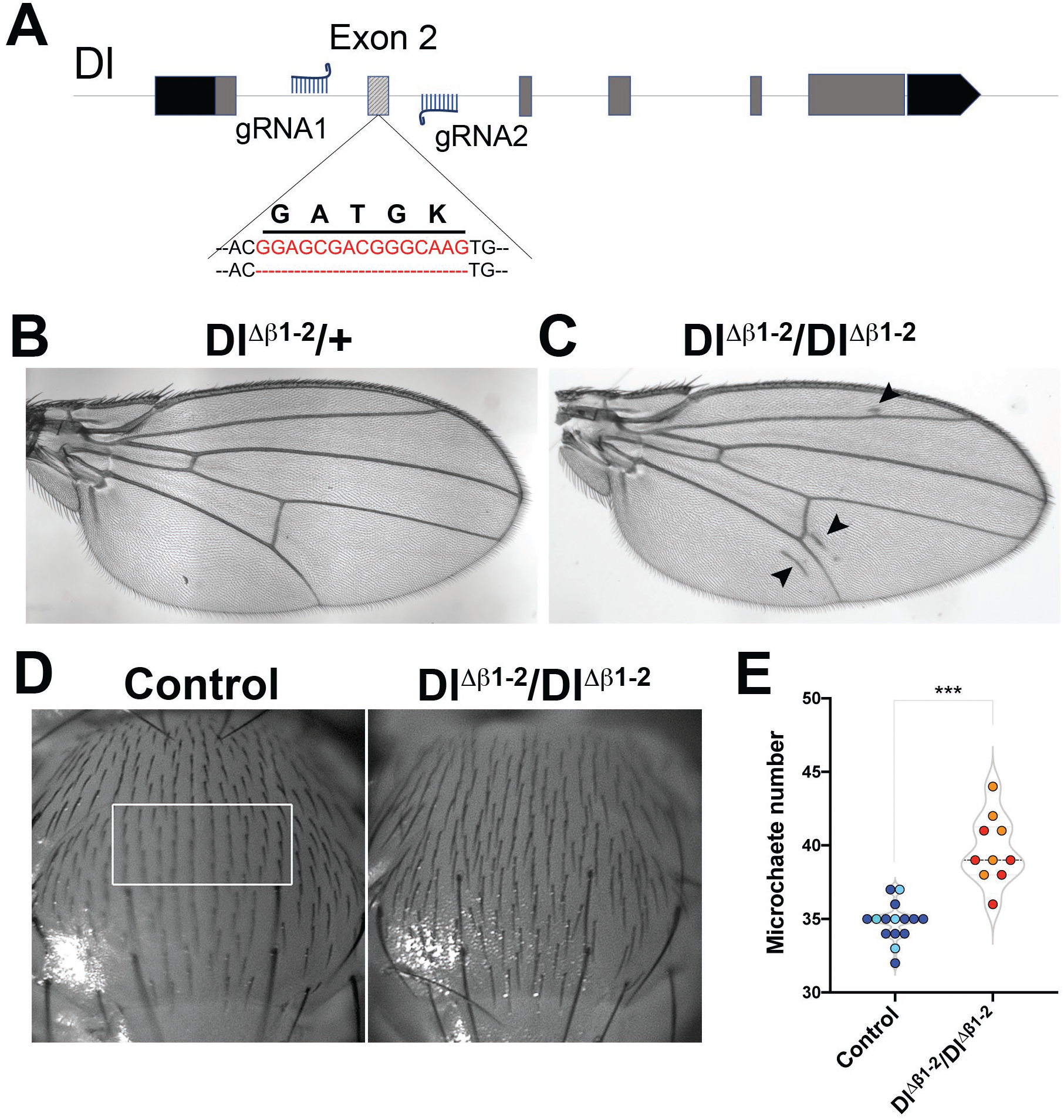
Dl β1-2 loop mutant generated by genome editing. (**A**) Two gRNAs flanking the *Dl* Exon 2 were used to replace the exon with a modified version where 5 amino-acids in the β1-2 loop were removed. Red lettering highlights the genomic sequence of the β1-2 loop. **(B-C)** Adult wings from *Dl^Δβ1-2^* flies. No defects are detected in wings from *Dl^Δβ1-2^/+* (B), Homozygous *Dl^Δβ1-2^/ Dl^Δβ1-2^* have extra vein tissue near L5 and uneven L2 veins (arrowheads; C). **(D)** Microchaetae are arranged in rows on the thorax of control (yw) flies; these become disordered and more dense in *Dl^Δβ1-2^/Dl^Δβ1-2^*. **(E)** Number of microchaetes per central area (as depicted by white rectangle in D) in the indicated genotypes, ***P<0.0001 (Unpaired t-test). Light, dark shading indicates data points from two independent replicates.

Severe loss of Delta function, as with *Dl^ΔExon2^*, results in lethality. In contrast *Dl^Δβ1-2^* homozygotes were viable. Nevertheless *Dl^Δβ1-2^* adult flies exhibited several visible phenotypes. Firstly, they had ectopic wing-vein material, with extra vein tissue detected around L2, L5 and the posterior cross-vein (Fig. 2C, arrowheads). Secondly, they had abnormal spacing of the microchaetae on the thorax (Fig. 2D-E). Both the venation and microchaetae defects are consistent with altered Notch pathway activity (Heitzler & Simpson, 1991; Vässin & Campos-Ortega, 1987), suggesting that localised mutations affecting the β1-2 loop impair the function of the Delta ligand.

The defects produced by *Dl^Δβ1-2^* were relatively mild, and there were no disruptions to the wing margin (e.g. notching). In agreement, expression of genes *cut* and *deadpan* that require high levels of Notch signaling at the d/v boundary was not disrupted in patches of *Dl^Δβ1-2^* mutant cells (Fig. EV3). Likewise, *Ser^Δβ1-2^* had normal wings (Fig. EV2B) but exhibited mild abnormalities associated with ectopic pigmentation of joints, that could also be indicative of compromised signalling. Together the data indicate that the specific deletion within the β1-2 loop has a detectable effect on Notch ligand functions.

### Ligand β1-2 loop mutants exhibit reduced activity

To further probe the consequences from the mutations in the C2 domain β1-2 loop, *Dl^Δβ1-2^* was combined in trans with previously characterised deletions (*Df(3R)Dl^Fx3^*) and loss of function (e.g. *Dl^rev10^*) *Dl* alleles. When heterozygous, the strong *Dl* alleles exhibit a robust and consistent wing vein phenotype, with “deltas” formed by extra vein material along several of the veins (Fig. 3A-C – left panel). In combination with *Dl^Δβ1-2^* this phenotype was strongly enhanced, so that more of the veins were affected and they became uneven and thickened (Fig. 3A-D – right panel). The enhancement of vein defects by *Dl^Δβ1-2^* occurred in combinations with all *Dl* alleles tested. Likewise, *Ser^Δβ1-2^* had a similar effect. Full Notch activity in the wing veins also requires Serrate, as revealed by chromosomes carrying mutations in both *Dl* and *Ser*, which have more severe phenotypes than *Dl* mutations alone despite the fact that *Ser/+* flies have normal veins (Fig. EV2B). Combining *Ser^Δβ1-2^* allele with this double mutant chromosome, enhanced the thickening of veins in a similar manner to *Dl^Δβ1-2^* (Fig EV2C-D). The enhanced vein phenotypes indicate that deletions within the β1-2 loop of the C2 domain compromise ligand activity.

**Figure 3.**
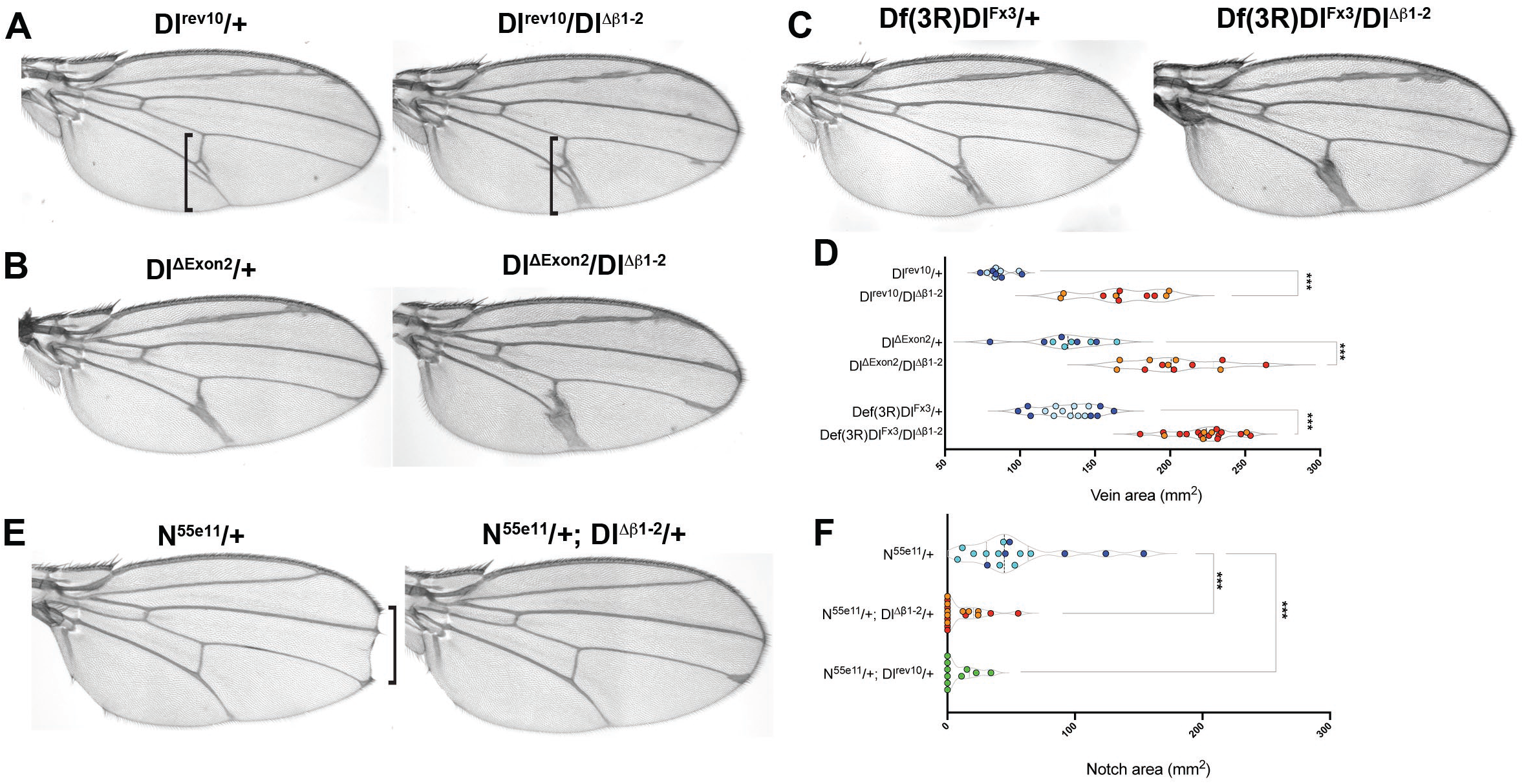
*Dl^Δβ1-2^* enhances vein thickening from loss of function Dl alleles and suppresses *Notch* phenotype. (**A-C**) Representative images of adult females wings in combinations of *Dl^Δβ1-2^* with loss of function *Delta* alleles. In combinations with *Dl^rev10^* **(A),***Dl^ΔExon2^* **(B),** or *Df(3R)Dl^Fx3^* **(C),** vein thickening is strongly enhanced (right panels) compared to heterozygous mutants alone (left panels). Square brackets indicate the regions used for vein thickness quantification. **(D)** Quantification of wing-vein thickness in females of the indicated genotypes. (**E**) Representative images of adult female wings demonstrate that *Dl^Δβ1-2^* rescues the wing notching phenotype, caused by a Notch loss of function allele (*N^55e11^*). **(F)** Quantification of wing notching in females of the indicated genotypes, *Dl^Δβ1-2^* rescues notching in a similar manner to *Dl^rev10^*; ***, P<0.0001(Unpaired t-test). Light, dark shading indicates data points from independent genetic crosses.

One unusual feature of the Notch pathway is that the ligand and receptor molecules can interact together in cis, when they are present on the same cell surface (De Celis & Bray, 1997). This cis-interaction is inhibitory and may be important to set a threshold that ensures a sharp response (Sprinzak *et al*, 2010). One manifestation of this balance is that the phenotypes produced by reduced *Notch* function are supressed when combined with a *Delta* loss of function allele. Notch heterozygous females have a characteristic wing-notching phenotype. When combined with *Dl^Δβ1-2^* the wing notching phenotype was suppressed to a similar extent as with a classic *Delta* allele, suggesting that cis-interactions are also modified.

Similar results were obtained with another Notch-dependent process, the spacing between the sensory organs, microchaetae, on the notum. In the absence of Notch signalling an excess of sensory organ precursors are formed due to failure in lateral inhibition (Cohen *et al*, 2010; Heitzler & Simpson, 1991; De Joussineau *et al*, 2003; Sjöqvist & Andersson, 2017). Milder defects in Notch signalling lead to irregular and reduced spacing between the sensory organ precursors with the consequence that there is an increase in the number of microchaete on the adult notum as seen in flies heterozygous for a deletion of Delta (e.g. *Df(3R)Dl^Fx3^/+*; Fig. EV4A-B,E). As noted above *Dl^Δβ1-2^* homozygous flies had an increased density of microchaetae compared to wild-type (Fig. 2D-E and EV4C,E) and in combination with strong Delta alleles, *Dl^Δβ1-2^* led to a further increase in microchaetae numbers (Fig. EV4D-E; *Df(3R)Dl^Fx3^/Dl^Δβ1-2^*). Thus, as with the vein formation, the defects in microchaetae spacing indicate a reduced signaling potential for ligands with a shortened β1-2 loop, despite the fact that this change should not disrupt binding to the receptor per se (see Fig. 1E).

### *Dl^Δβ1-2^* has compromised Notch signalling in photoreceptor fate decisions

Flies homozygous for *Dl^Δβ1-2^* also had mild roughening of the eyes. Notch activity is required at several stages in the development of the photoreceptors, including in the specification of R4 and R7 photoreceptors. The sequential differentiation of the eight neuronal photoreceptors (R cells) is initiated when a wave of differentiation (called Morphogenetic Furrow or MF) spreads from the posterior to the anterior region of the eye imaginal disc ((Şahin & Çelik, 2013); Fig. 4A). Notch activity in one cell of the five-cell cluster specifies R4 cell fate and can be detected by the expression of E*(spl)mδ0.5-lacZ*, containing the Notch responsive *E(spl)mδ* enhancer ((Cooper & Bray, 1999); Fig. 4A,B). Reducing the levels of Delta, as seen in Delta heterozygotes *Df(3R)Dl^Fx3^*/+, led to more variable expression of E(spl)mδ0.5 (Fig. 4B). This was further enhanced in combination with *Dl^Δβ1-2^*, so that many of the ommatidia exhibited very low levels of expression (Fig. 4B-C). No similar reduction occurred with *Dl^Δβ1-2^* heterozygotes (Fig. 4B) nor clones of *Dl^Δβ1-2^* homozygous mutant cells (Fig. EV5) arguing that the decrease in activity in these conditions is not below the threshold needed for E(spl)mδ0.5 activation. Nevertheless, the fact that the *Dl^Δβ1-2^* enhances the phenotype from the Delta deletion is consistent with it being compromised for productive Notch signalling.

**Figure 4.**
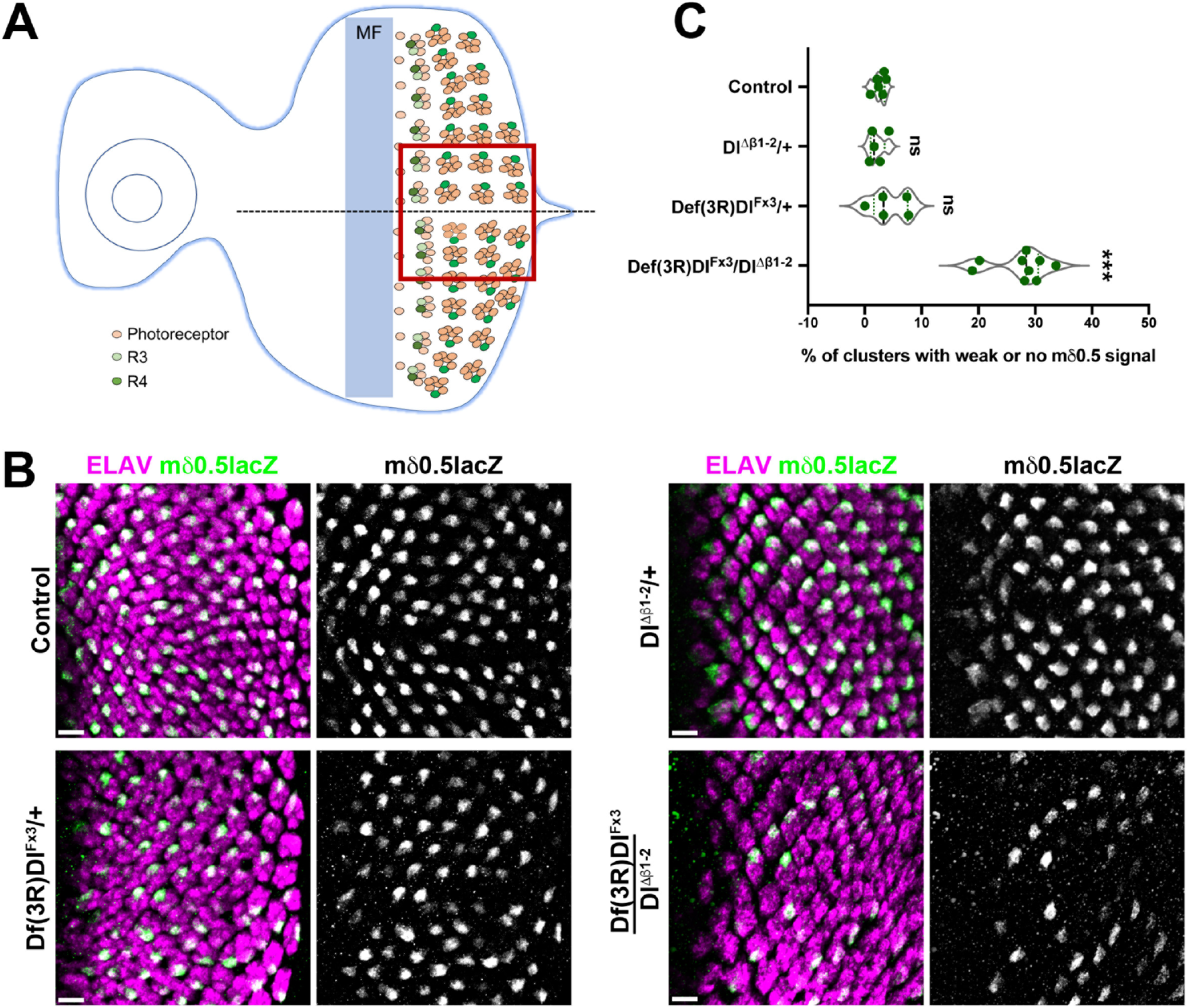
*Dl^Δβ1-2^* has compromised Notch response in photoreceptor fate decisions. (**A**) Schematic representation of Notch reporter *E(spl)mδ0.5* expression during photoreceptor differentiation. Expression is initiated in R3 and R4 of the 5-cell pre-cluster and becomes restricted to R4 as Notch activity resolves. Light orange indicates photoreceptors with R3 in light green and R4 in dark green. MF marks the morphogenetic furrow, boxed region indicates the region shown in B. (**B)** Equatorial region of eye imaginal discs where *E(spl)mδ0.5* expression (green) becomes restricted to a single photoreceptor in each cluster (magenta), as detected in control and *Dl^Δβ1-2^*/+ discs (top panels). In *Df(3R)Dl^Fx3^* /+ and *Dl^Δβ1-2^/Df(3R)Dl^Fx3^* discs (bottom panels) E(spl)mδ0.5 expression is reduced (*Df(3R)Dl^Fx3^* /+) or absent from several clusters (*Dl^Δβ1-2^/Df(3R)Dl^Fx3^*) indicative of reduced Notch signalling. **(C)** Proportion of photoreceptors clusters that fail to express the *E(spl)mδ0.5* reporter in the indicated genotypes. ns, no significant difference, *** P<0.0001 (one-way ANOVA). Scale bars correspond to 10 μm.

### C2 Domain β1-2 loop mutation does not impair Delta trafficking

Our results indicate that the β1-2 loop region of Delta C2 domain is required for full functionality. To investigate whether this involves a change in the localization or trafficking of Delta we generated mutant clones in the wing disc, a tissue where the expression and localization of the ligand is well characterised. In late third instar stages, the expression of Delta is particularly enriched in two stripes flanking the DV boundary and in longitudinal stripes that prefigure the prospective wing veins (Fig. 5A,A’). In all regions of the disc, Dl^Δβ1-2^ exhibited normal expression levels and it appeared to be localized at the apical membranes, at similar levels to wild-type Delta.

**Figure 5.**
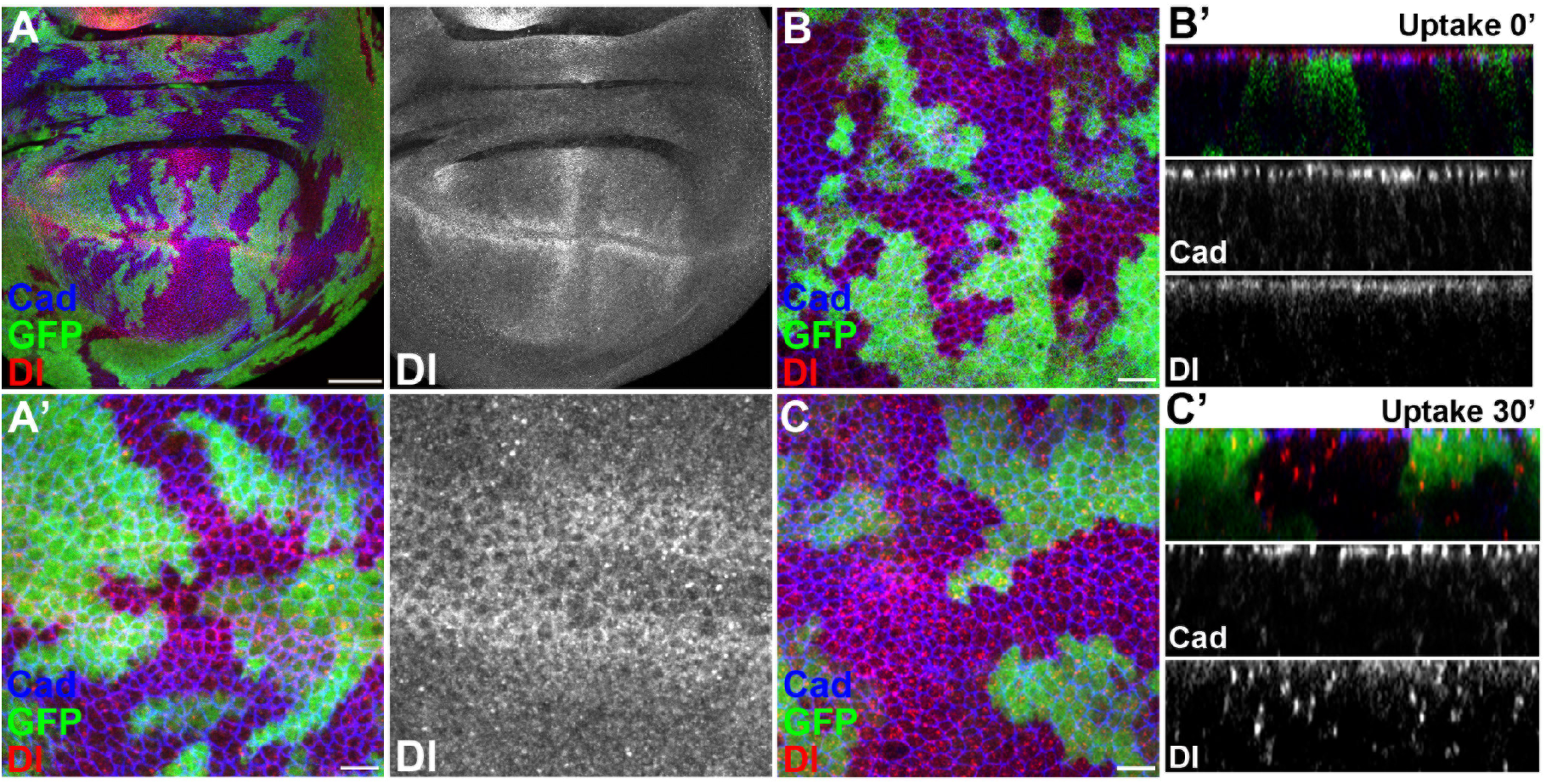
*Dl^Δβ1-2^* exhibits normal sub-cellular localisation. (**A**) Apical view of wing imaginal disc with homozygous *Dl^Δβ1-2^* clones (GFP negative) stained for Dl (red) and Cadherin (blue). (**A’**) Z-projection of apical layers spanning a *Dl^Δβ1-2^* clone (GFP negative) located at the DV boundary. No change in apical localisation of Dl (red) and Cadherin (blue) is detected. **(B)** Uptake assay at t=0. After exposure to extracellular anti-Dl antibody, Dl protein (red) is detected at similar levels apical to Cadherin (blue) in wild-type (GFP) and *Dl^Δβ1-2^* mutant tissue (GFP negative). **(B’)** Cross-sectional view of **B**, Dl protein (red) is present apically relative to Cadherin (Blue) which marks adherens junctions. **(C)** Uptake assay after 30 minutes, internalised anti-Dl (red) enters the endocytic route and in the cross sectional view **(C’)** can similarly be detected as puncta along the cell axis in wild-type (GFP) and homozygous *Dl^Δβ1-2^* (GFP negative) tissue.Scale bars: A, 50 μm; A’ B and C,10 μm.

To confirm that the mutant protein was present on the surface we performed an antibody uptake assay (Le Borgne & Schweisguth, 2003). Wing imaginal discs were incubated *ex vivo* with an anti-Dl antibody recognizing the extracellular domain at 4°C. Excess antibody was then washed away, and the tissues transferred to a permissive temperature (25°C) for 0 or 30 minutes so that the membrane localization, uptake and trafficking of bound antibody could be measured (Gomez-Lamarca *et al*, 2015). At zero minutes when antibody was bound to Delta on the cell-surface, similar levels were detected in control regions and in *Dl^Δβ1-2^* mutant clones (Fig. 5D-D’’), indicating that the mutant protein was present on the cell surface. When endocytosis was allowed to proceed for 30 minutes, antibody bound Delta accumulated in puncta throughout the epithelial cells in both wild-type and Dl^Δβ1-2^ tissue. The uptake assays confirm therefore that the mutated protein reaches the cell surface normally and that its trafficking following endocytic uptake is unaltered.

## DISCUSSION

C2 domain phospholipid binding properties are essential for membrane-targeting of many intracellular proteins. Notch ligands are unusual in having an extracellular N-terminal C2 domain (Chillakuri *et al*, 2013; Kershaw *et al*, 2015). This structure is present in all the human Notch ligands and retains the capacity to interact with liposomes (Suckling *et al*, 2017). Here, we showed that *Drosophila* Delta and Serrate also contain a globular C2 domain that confers the ability to bind to phospholipid-containing liposomes *in vitro*. The C2 domain structures are highly conserved, differing only in the length and orientation of several loops. A deletion mutation affecting one of these, a loop between the β1 and β2 strands of the C2 domain core, was sufficient to compromise liposome binding. This loop might therefore help to generate a “pocket” capable of interacting with a specific type of lipid, e.g. phospholipid/glycosphingolipid and in this way influence productive Notch signalling.

A subset of human *Jagged1* mutations that affect the loops at the apex of the C2 domain are associated with extrahepatic biliary atresia suggesting these regions are important for tuning the Notch signal in physiological contexts (Kohsaka *et al*, 2002; Suckling *et al*, 2017). Our results, from CRISPR engineering β1-2 loop mutations in Drosophila Delta and Serrate, support the conserved functional importance of the C2-domain loops. The mutated Delta exhibited reduced signaling activity in several different developmental contexts. The compromised signaling was most evident in genetic combinations with a strong loss of function allele or deletion of the locus, and was manifest by enhanced vein thickening, extra sensory bristles and reduced signalling during photoreceptor fate choice although there were no overt effects at the dorsal-ventral boundary. All of the processes affected involve highly dynamic signaling and are sensitive to subtle changes in signaling as evident from the defects in animals with reduced dosage of wild-type Delta (*Df(3R)Dl^Fx3^*/+). These results are consistent with the model that C2 domain loop regions are important for fine-tuning the Notch signal (Fig. 6), as suggested by *in vitro* results coupling C2 domain lipid binding to Notch binding and with the changes in Notch activation seen with EHBA and related loop variants (Suckling *et al*, 2017).

**Figure 6.**
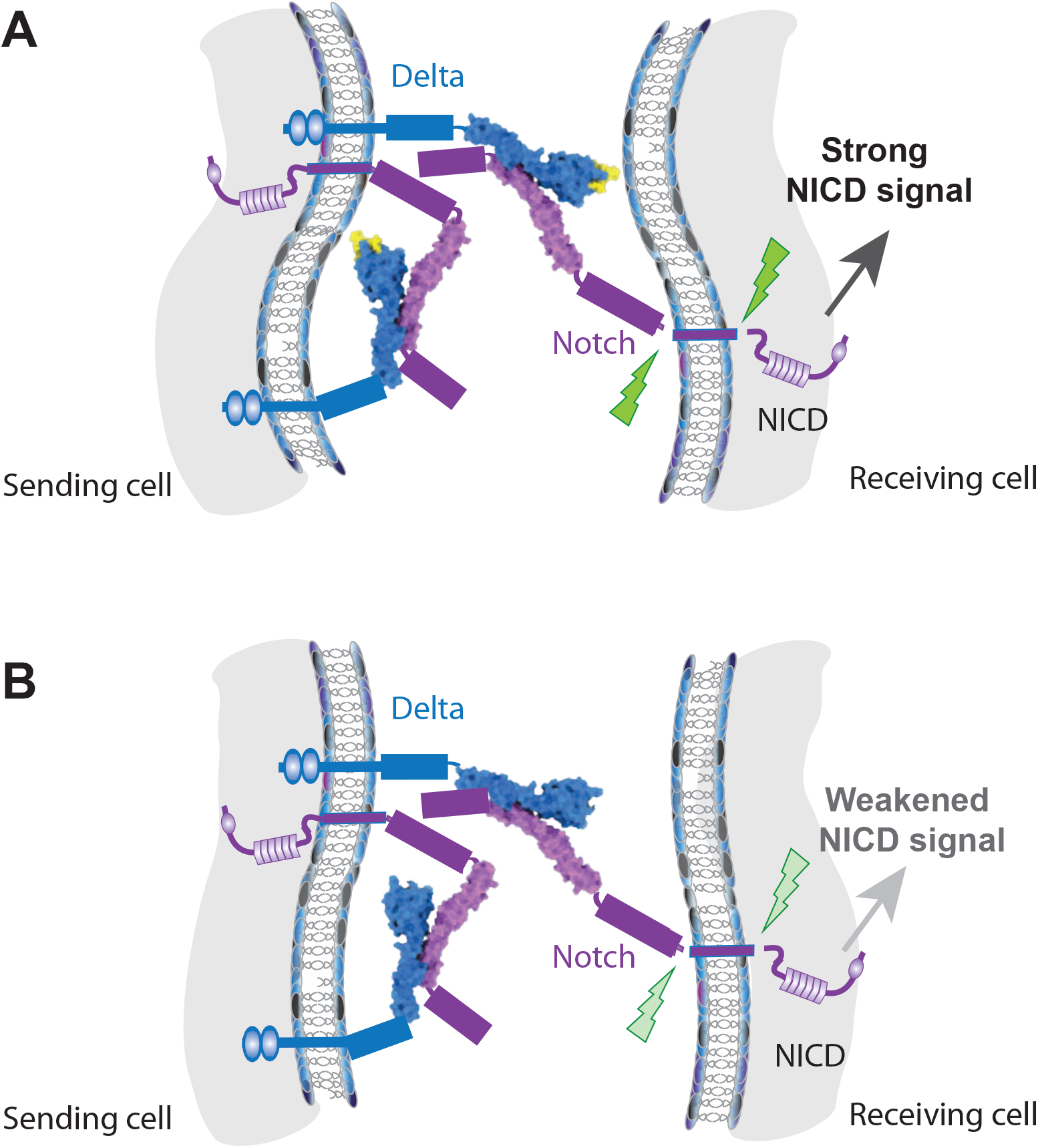
Schematic summarizing roles for ligand β1-2 loop. (**A, B**) The C2, DSL domains and N-terminal EGF repeat in Delta (cyan) and the ligand binding region (EGF11-13) of Notch (magenta) are renditions from the structures obtained (Figure 1), other regions of the molecules are represented not to scale. The amino acids in the β1-2 loop of Delta are highlighted in yellow (A). **(A)** The interaction of Delta (cyan) in *trans* with the Notch receptor (magenta) is augmented by the C2 domain, possibly through contacts of the β1-2 loop (yellow), with the “receiving” cell membrane, to yield highest levels of signalling (black arrow; green indicates ligand induced cleavages). Phospholipid contacts from β1-2 loop in the same cell could also influence cis-interactions between Delta and Notch in the same cell. **(B)** A deletion of 5 amino acids within C2 domain β1-2 loop (no yellow) disrupts phospholipid interactions but does not prevent Delta from interacting with Notch. Activation of Notch signalling is weakened (grey arrow) and phenotypes from transheterozygous combinations suggest that cis-interactions between Delta and Notch are also modulated.

There are several models for how C2 domain-mediated membrane interactions might impact on signalling. One possibility is that the spatial or temporal residence of Delta in the membrane may be affected by the C2 domain interactions. Evidence suggests that relative pools of the receptor and ligands, rather than absolute concentrations, are important for refining signaling outcomes due to the balance between cis-inhibition and trans-activation (Sprinzak *et al*, 2010). Models based on this relationship inferred that intrinsic noise would cause the width of the vein to become irregular, one characteristic of the phenotype produced from *Dl^Δβ1-2^*. Loss of interaction with certain types of lipids might bias how the ligand interacts with the receptor in favour of inhibition, producing a generalized reduction in signaling, despite there being similar amounts of proteins on the cell surface. For example, C2 domain interactions could affect the length of time the ligand is diffusing in the membrane and consequently fine-tune the levels of signalling (Khait *et al*, 2016).

In summary, our structure-guided approach to make defined changes in the endogenous ligands has demonstrated the *in vivo* relevance of C2 domain loops for full activity in the physiological setting (Fig. 6). This approach has uncovered subtle functional requirements that would unlikely be detected using *in vitro* or *in vivo* methods alone, highlighting the importance of using interdisciplinary methods to fully elucidate function.

## MATERIALS AND METHODS

### Protein Expression, Crystallisation and Structure Determination

Codon optimized open reading frames for constructs (synthesized by GeneArt®, Life Technologies Ltd., Paisley,UK), with recommended BiP signal peptide (for secretion), were subcloned into expression vector pEXS2-2 (Expres2ion® Biotechnologies, Horsholm, Denmark) using *Eco*RI and *Not*I restriction sites (see Table EV1 for primer sequences). Each construct was expressed as a monomer with a C-terminal 8xHis tag to facilitate purification. Purification was as described in Suckling et al.(Suckling *et al*, 2017). *Drosophila* (d) Delta and Serrate NE3 (C2 domain-DSL, EGF1, EGF2, EGF3) constructs were used for liposome- and Notch-binding assays. NE3 (residues 1-332 Delta; 1-388 Serrate), NE2 (residues 1-293 Delta, residues 1-349 Serrate), NE1(1-259 Delta, 1-314 Serrate) constructs were used to set up preliminary crystal trials. Delta NE1 and Serrate NE2 constructs produced best diffracting crystals. NotchEGF11-13 was produced using the same expression system, and the purified cleaved form used for crystallization.

The Notch receptor construct was concentrated to 24.2mg/ml in a buffer A (10mM Tris pH 7.5, 150mM NaCl, 10mMCaCl_2_) and crystallised by the sitting drop method from 200nl + 200nl drops with mother liquor 0.1M MES pH 6.5, 1.8M Ammonium sulphate, 0.01M cobalt chloride. Crystals were cryoprotected by addition of 25% ethylene glycol and data collected at the European Synchrotron Radiation Facility, beamline ID29. Serrate was concentrated to 15.8 mg/ml in buffer A and crystallised by the sitting drop method from 200nl + 200nl drops with mother liquor 0.1M imidazole malate pH 7, 25% PEG4K and cryoprotected by addition of 25% ethylene glycol, 20 mM CaCl_2_. Data were collected at Diamond Light Source, beamline I02.Delta was concentrated to 17.7 mg/ml and crystallised by the sitting drop method from 200nl + 200nl drops with mother liquor 0.1M Tris pH 8.5, 0.2M MgCl2, 30% PEG4K and cryoprotected with 25% glycerol, 20mM CaCl2. Data were collected at Diamond Light Source, beamline I04.

All structures were solved by molecular replacement using separated domains from the human homologues using program PHASER (McCoy *et al*, 2007) from program suite CCP4 (The CCP4 suite: programs for protein crystallography., 1994) rebuilt using BUCCANEER (Cowtan, 2006) and COOT (Emsley *et al*, 2010) and refined in PHENIX (Liebschner *et al*, 2019). For all constructs data processing and model statistics are described in Table 1. Coordinates and data are deposited in the Protein Data Bank with accession codes 7ALJ, 7ALT, 7ALK for Notch, Serrate and Delta respectively.

### Liposome and Notch Binding assays

Liposome binding assays were carried out as described in Suckling, Korona et al (Suckling *et al*, 2017) using purified Delta/Serrate variants and liposomes comprising PC:PS:PE-fluoroscein in a 80:15:5 ratio. Liposomes were prepared as described in Chillakuri *et al* (Chillakuri *et al*, 2013). Notch binding assays were carried out as described in (Suckling *et al*, 2017) using purified Delta variants and Notch EGF 11-13. The negative control Delta F204A variant reduces Notch/ligand binding at Site 2, by altering a key residue within the ligand DSL domain Notch-binding loop.

### *Drosophila melanogaster* strains and genetics

All *Drosophila melanogaster* stocks were grown on standard medium at 25°C. Alleles are as described in Flybase(Thurmond *et al*, 2019) and in particular the following were used to sensitize the genetic background: *Dl^rev10^* (Heitzler & Simpson, 1991), *Dl^rev10^,Ser^Rx106^* (Thomas *et al*, 1991), *Df(3R)Dl^Fx3^* (Vässin & Campos-Ortega, 1987), *N^55e11^* (#BL28813). *Dl^Δβ1-2^* clones were generated using FRT mediated recombination (Xu & Rubin, 1993) – recombination was promoted by heat-shock of 1h at 37°C 72h prior to dissection and analysis. The *E(spl)mδ0.5* reporter was used for analysis of R3/R4 determination in eye imaginal discs, (Cooper & Bray, 1999).

### Generation of β1-2 loop Notch Ligands mutants using CRISPR/Cas9

Lines were generated by CRISPR-mediated homology repair (HR) strategy. As described in Fig 3 and FigEV2, two guideRNAs were designed to flank the target exon coding the β1-2 loop (see Table EV1 for primer sequences) and cloned into the guide RNA expression pCFD4 vector (Addgene #49411). The exon of interest and homology arms were cloned into donor template plasmid pHD-ScarlessDsRed (Addgene # 64703) using the Gibson Assembly Protocol (see Table EV1 for primer sequences). Modifications to the exons were made using standard mutagenesis and PCR amplification prior to the co-injection of the guide RNAs and the donor template constructs into *nos-Cas9* (#BL54591) embryos. Modifications included: 1) deletion of 15bp within β1-2 loop of Dl (*Dl^Δβ1-2^*); 2) deletion of 15bp within β1-2 loop from *Ser* Exon 3 (*Ser^Δβ1-2^*); 3) deletion of region between the two gRNAs (*Dl^ΔExon2^*). Engineered flies were identified by expression of DsRed in the eyes and verified by genomic PCR sequencing. The transposable element containing the DsRed was removed subsequently by crossing to flies carrying PiggyBac Transposase (#BL32070).

### Immunostainings

The following primary antibodies were used for Immunofluorescence staining: Goat anti-GFP (1:200, Abcam, ab6673), Mouse anti-Cut (1:20, Developmental studies hybridomw bank (DSHB)), Rat anti-DE-Cad2 (1:200, DSHB), Mouse anti-Delta (1:30, DSHB), Guinea pig anti-Delta (1:2000, a gift from Mark Muskavitch, (Huppert *et al*, 1997)), guinea pig Anti-Dpn (1:2000, a gift from Christos Delidakis), Mouse anti-NECD (1:50, DSHB), Rat anti-ELAV (1:200, DSHB), Mouse anti-β-Gal (1:1000, Promega, Z378A). Uptake assay was performed as described previously (Gomez-Lamarca *et al*, 2015).

### Adult tissues analysis

For the analysis of the adult fly wings, female flies were collected in 70% Ethanol for 2 hours, rehydrated in PBS and one wing per fly was isolated and mounted in a 50% glycerol solution. To analyse the microchaete number, flies were collected in 70% Ethanol for 2 hours, rehydrated in PBS 1x and mounted on apple juice agar plates for imaging.

### Image and statistical analysis

Immunostaining samples were imaged with Leica TCS SP8 microscopes (CAIC, University of Cambridge) at 40X magnification and 512/512 or 1024/1024 pixel resolutions. Images of the adult wings were taken using a Zeiss Axiophot microscope and images of the adult Notum were taken using the Leica MZ10F coupled with a camera Leica DFC3000G. ImageJ software was used to analyse images and polygon tool was used to measure the vein area on the region limited by the CV2, L4 and L5 veins on adult wings. The measurement of the wing notching was done by determining the tissue missing with the polygon tool after superimposing wings of the described genotypes with the reference wild type wing. The number of microchaete was assayed using a fixed area as reference on the Notum, as depicted by the white box on figure EV4A. For the analysis of the Dl and Notch trafficking in *Dl^Δβ1-2^* mutant clones, a projection of 3-cell diameter was performed after re-slicing the images into the XZY axis in ImageJ software.

Statistics were calculated with GraphPad Prism. Comparisons between two groups were performed with a two-tailed unpaired t-test. Statistical differences among various groups were assessed with ordinary one-way ANOVA by comparison to the mean of the control column. ns indicates no significant difference; * P<0.1; ** P<0.001; ***P<0.0001;

## Data Availability

Coordinates and data have been deposited in the Protein Data Bank with accession codes 7ALJ, 7ALT, 7ALK for Notch, Serrate and Delta respectively.

## Acknowledgments

We thank Mark Muskavitch, Christos Delidakis, the Bloomington Stock Center, the VDRC Stock Center and the Developmental Studies Hybridoma Bank for *Drosophila* strains and antibodies. We thank Kat Millen for the technical help. We thank other members of SJB lab for valuable discussion. This work was funded by project grants from BBSRC (BB/P006175/1 to SJB) and MRC (MR/L001187/1 to SML and PH; MR-R009317/1 to PH) and by a Wellcome Trust investigator Award (100298 to SML)

## Author contributions

T.M and S.J.B designed the *in vivo* experiments. T.M performed the *in vivo* experiments. T.M and S.J.B analysed the data. P.H and S.M.L designed the *in vitro* experiments. B.K, Y.M and R.S purified ligand and receptor proteins. R.S and S.J performed structure determination. Y.M performed the Notch-binding experiments. B.K performed liposome-binding assays. B.K, Y.M, R.S, P.H and S.M.L analysed data. T.M, P.H, S.M.L and S.J.B wrote the manuscript.

---

**Table EV1.**
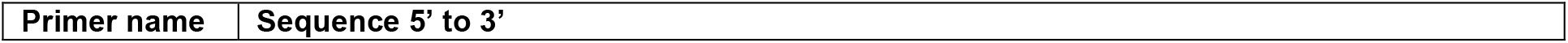
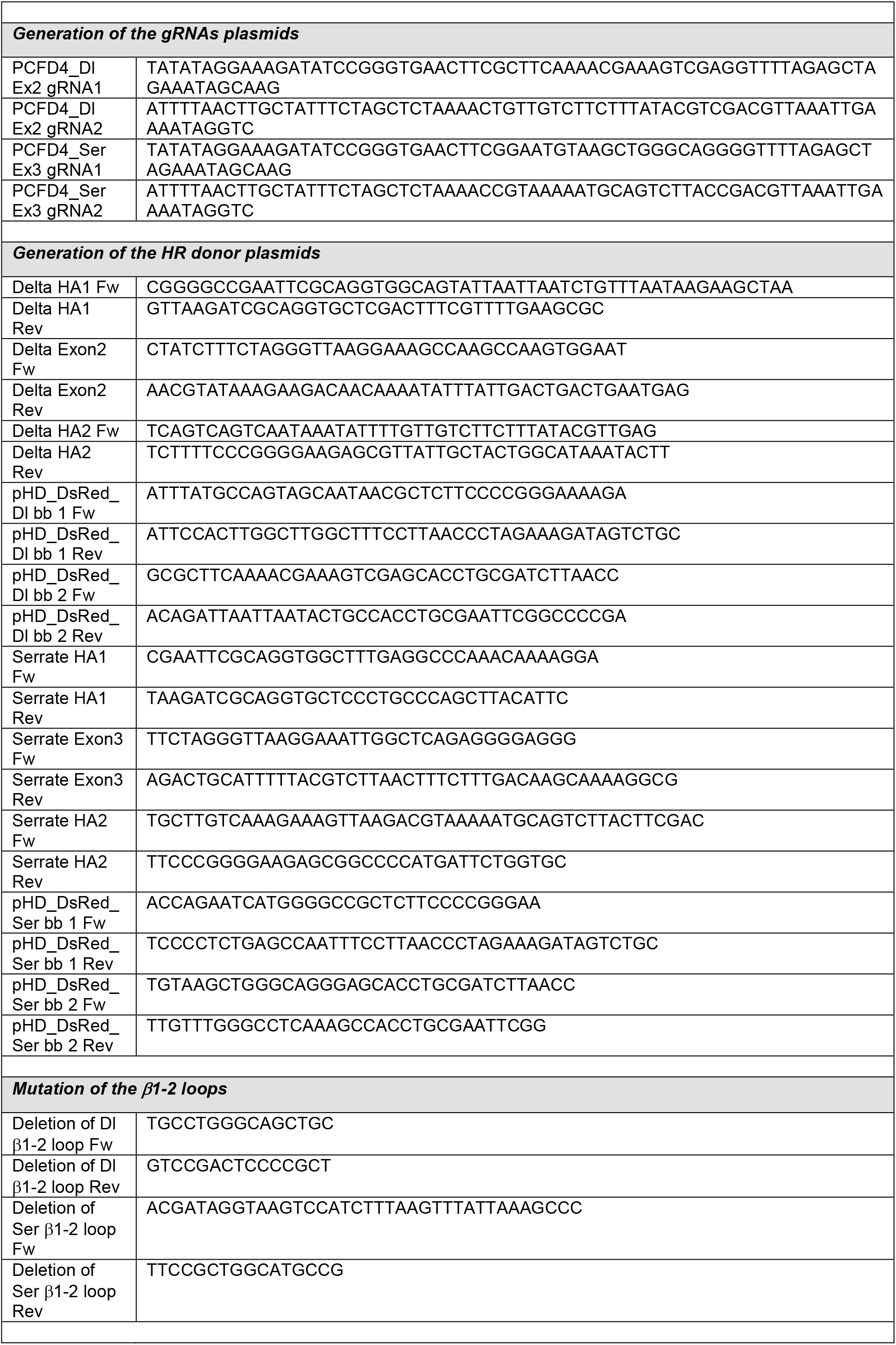
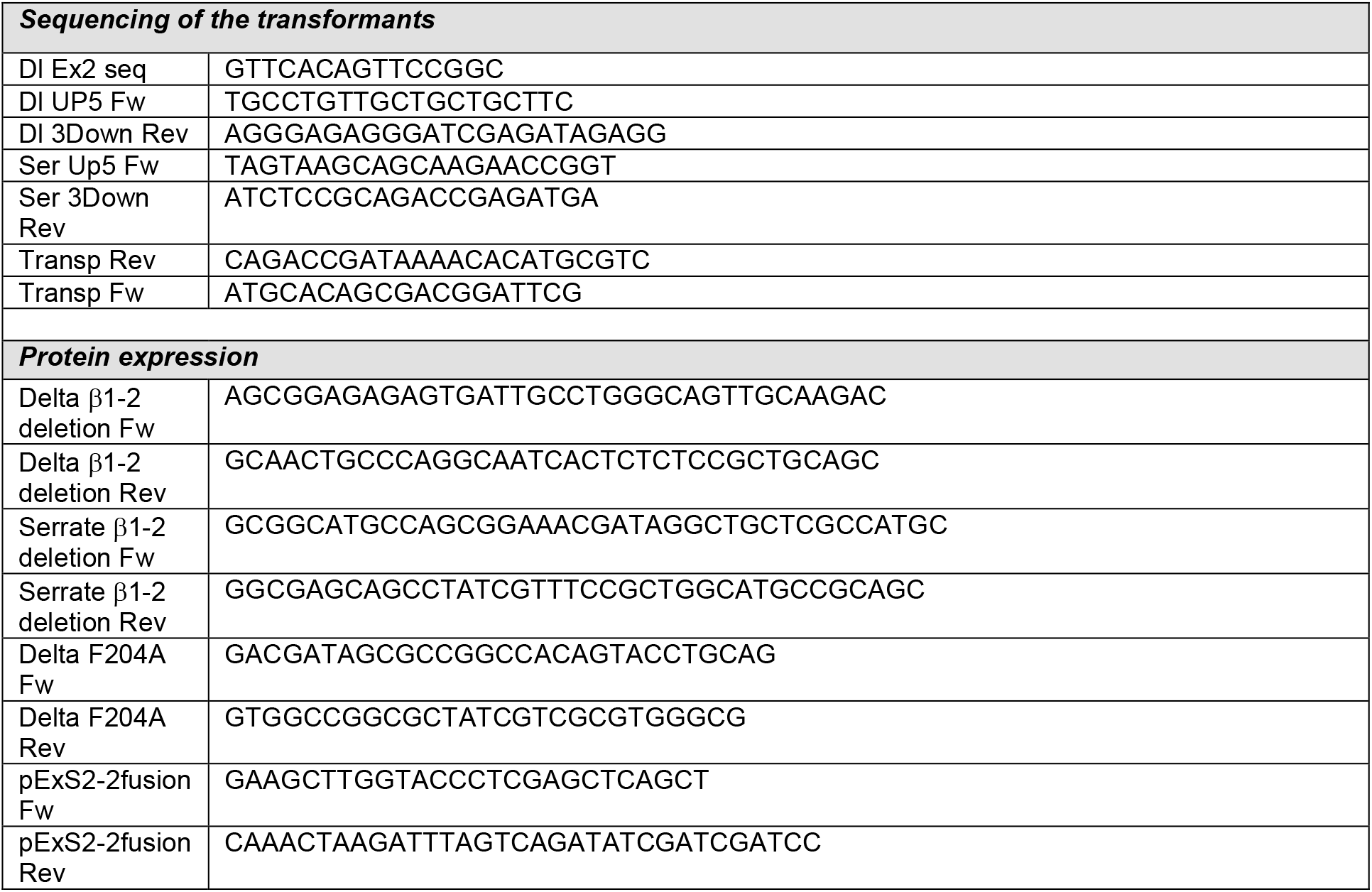
Primers list.

